# Variations in soil nutrient dynamics and bacterial communities in long-term tea monoculture production systems

**DOI:** 10.1101/2021.04.25.441296

**Authors:** Heng Gui, Lichao Fan, Donghui Wang, Peng Yan, Xin Li, Yinghua Pang, Liping Zhang, Kazem Zamanian, Jianchu Xu, Wenyan Han

**Author notes:** Corresponding Authors: Lichao Fan, Wenyan Han.

## Abstract

Long-term monoculture agriculture systems could lead to soil degradation and yield decline. The ways in which soil microbiotas interact with one another, particularly in response to long-term tea monoculture systems are currently unclear. In this study, through the comparison of three independent tea plantations across eastern China composed of varying stand ages (from 3 years to 90 years after conversion from forest), we found that long-term tea monoculture led to significant increases in soil total organic carbon (TOC) and microbial nitrogen (MBN). Additionally, the structure, function and co-occurrence network of soil microbial communities were investigated by pyrosequencing 16S rRNA genes. The pyrosequencing analysis revealed that structures and functions of soil bacterial communities were significantly affected by different stand ages of tea plantations, but sampling sites and land-use conversion (from forest to tea plantation) still outcompeted stand age to control the diversity and structure of soil bacterial communities. Further RDA analysis revealed that the C and N availability improvement in tea plantation soils led to variation of structure and function in soil microbial communities. Moreover, co-occurrence network analysis of soil bacterial communities also demonstrated that interactions among soil bacteria taxa were strengthened with the increasing stand age of respective tea stands. Overall, this study provides a comprehensive understanding of the impact of long-term monoculture stand age on soil nutrient dynamics and bacterial communities in tea production.

## 1. Introduction

Soil microbial communities play an indispensable role in maintaining soil health and nutrient cycling (Bünemann et al., 2018; Torsvik et al., 1996) and are strongly affected by environmental factors (e.g., soil pH, soil texture, nutrients availability) (Geisseler and Scow, 2014; Girvan et al., 2003; Lauber et al., 2009), plant species (El Zahar Haichar et al., 2008), land management (Rodrigues et al., 2013), and locations (Decaёns, 2010; Fierer and Jackson, 2006; Martiny et al., 2006). Among these factors, land use change is the most impactful factor via that human disturb soil environmental conditions, thereby altering the structure, diversity, and biomass of bacterial communities (Da C Jesus et al., 2009). The type of vegetation planting (El Zahar Haichar et al., 2008) and management (Geisseler and Scow, 2014) along a chronosequence (i.e. land-use duration) after land use conversion play important roles in controlling the variation of soil bacterial communities. This is because of reshaping the soil structure, accumulation of soil nutrients and hazardous substances.

Microorganisms usually form complex interactive networks in which interactions among members are essential for community assembly and ecosystem functions (Deng et al., 2016; Shi et al., 2016). Therefore, identifying and defining the interactions that occur among soil microorganisms are critical to understanding microbial diversity and functions (Banerjee et al., 2016). Network analysis of co-occurrence, as usually determines by correlations between abundances of microbial taxa provides a promising start for exploring the organization and dynamics of microbial interactions and niches (Barberán et al., 2012; Chen et al., 2019; Jiao et al., 2016). Exploring these microbial interactions, rather than those of simple richness and composition involved in soil environment, especially in agriculture soils, can provide important information on plant health and growth (Banerjee et al., 2016; Zhang et al., 2018). Recent advances in high-throughput sequencing approaches now enable us to apply network analyses to explore more information in the complex uncultivated soil microbial communities of agricultural land. For example to define network complexity or stability between microbial communities and environmental factors (Ma et al., 2020) or identifying potential keystone species (Fan et al., 2018) or geographical patterns at a continental scale (Zhang et al., 2018). However, there is little information about topological variation found in soil microbial co-occurrence interactions in long-term monoculture agricultural systems.

Tea (*Camellia sinensis* L.) is a perennial evergreen broad-leaved cash crop (Han et al., 2007), and tea is one of the top three consumed beverages in the world. Tea plantations are typically established by conversion from forest and with years (decades to centuries) of intensively managed cultivation practices (Han et al., 2007). Most tea plantations are distributed in subtropical areas, and over 3.06 million ha are established in China, a figure which is only increasing (Fan and Han, 2020). Soil degradation is one potential issue arising from long-term monoculture of tea plantations (Yan et al., 2018; Yan et al., 2020), though providing an accumulation of C and N (Fan and Han, 2020; Fan et al., 2015). To better understand the mechanisms undergirding soil nutrients-cycling network in tea plantations, that promotes soil fertility as well as production and quality of tea, it is of vital importance to document the influence of land-use change and long-term monoculture on soil microbiomes in tea plantations. Accordingly, increased attention has been concentrated on soil microbiomes in tea plantations, such as microbial community structure and microbial biomass as affected by the stand age of tea plantations (Wang et al., 2019). However, the relative importance of long-term monoculture systems and spatial variation on the structure and function of bacterial communities in tea plantations remains unclear.

Here, we analyzed soil bacteria communities via pyrosequencing analysis at varying stand ages (from 3 to 90 years) across tea plantations at three separate sites (Fig. S1) and their adjacent forests. Our study aimed to answer the following research questions:

i. How do stand age (time after land conversion) and the sites affect the structure and function of soil bacterial communities in tea plantations?
ii. Which possible environmental factors could lead to the changes in soil bacterial communities described in i.?
iii. How does stand age affect the interactions in soil bacterial communities in tea plantations based on co-occurrence network analysis?

## 2. Materials and methods

### 2.1. Experimental design and soil sampling

The sampling of tea plantation sites is shown in Fig. S1, and selected environmental information is provided in Table S1. In general, three tea plantations composed of tea stands with varying stand ages in Zhejiang Province, China were chosen for comparing changes in soil bacterial communities between different tea stand ages. The first one was located in Jingning County, Lishui City (JL), in which three tea stands aged 3 (Y3), 21 (Y21), and 43 (Y43) years were selected. The second was at the Tea Research Institute of the Chinese Academy of Agricultural Sciences (TRI), in which two tea stands aged 10 (Y10) and 90 (Y90) years were selected. The last one was located in Wenjiashan Village, Hangzhou City (HZ), in which three tea stands aged 13 (Y13), 50 (Y50), and 90 (Y90) years were selected. For these three different sites the annual mean temperature is 17 ° C, ranging from 1.7 ° C in January to 33.0 ° C in July. The annual mean precipitation is 1533 mm, with 74% of total rainfall occurring during the tea growing season from March to September. At JL and HZ sites, one forest (the land use prior to clearing and planting tea) adjacent to tea stand was selected for comparison. As described in Han et al. (2007), forest vegetation in both HZ and JL sites was dominated by *Cyclobalanopsis glauca* and *Quercus acutissima* Carri. The management for different tea stands at each tea plantation were similar. As described in our previous study (Han et al., 2007), 2250 kg ha^-1^ organic fertilizer (mainly rape seed cake containing 45% organic C, 4.6% N, 0.9% P, and 1.2% K) or 1500 kg ha^-1^ compound fertilizer containing approximate 8% N, 3.4% P, and 6.6% K was applied every September or October.

### 2.2. Soil sampling and treatments

For each tea stand at eachtea plantation, three 400 m^2^ plots of representative soil were randomly selected for soil collection. For each plot, eight random soil sub-samples at 0-20 cm depth were taken and mixed into one independent soil sample for later related analysis. Before soil sampling, the litter layer was removed at each plot. The soil was sieved using a 2 mm pore-size screen to remove plant roots, stones, and soil fauna. Each independent soil sample was divided into two parts. One part (50g) was stored at 4 ° C for later soil physicochemical analysis. The other part (10g) was stored at - 80 ° C for later DNA extraction.

### 2.3. Soil physicochemical properties analysis

Soil pH was determined by a combination of glass electrodes using a 1: 2.5 (w: v) ratio of soil to distilled water. Soil total C (TOC) and N (TON) were measured by LECO CNS Combustion Analyzer (LECO, CNS 2000, LECO Corporation, Michigan, USA) following manufacturer protocol. Soil microbial biomass C (MBC) and N (MBN) were determined following the fumigation-extraction method. Available phosphorus (AP) was extracted using hydrochloric acid and ammonium fluoride and determined using the molybdenum blue method. The concentration of exchangeable K (Exch. K), Ca (Exch. Ca), Mg (Exch. Mg), and Na (Exch. Na)were tested following hot block acid digestion protocol (Huang, Schulte et al. 1985).

### 2.4. Soil microbial DNA extraction and PCR amplification

Total DNA was extracted from about 0.5g of soil from each sample using the Mo Bio PowerSoil DNA isolation kit (Carlsbad, CA, USA) according to the manufacture’s instruction. After extraction, quality and concentration of DNA were tested using the NanoDrop ND 200 spectrophotometer (Thermo Scientific, USA). In accordance with the concentration, all DNA samples were diluted to 1 ng/uL before PCR amplification.

The V4 and V5 variable region of the bacterial 16S rRNA gene was amplified using the primers 515F (5’-CCATCTCATCCCTGCGTGTCTCCGAC-3’) and 907R (5’-CCTATCCCCTGTGTGCCTTGGCAGTC-3’). The polymerase chain reaction (PCR) amplification mixture was prepared with 1 μL purified DNA template (10 ng), 5 μL 10 × PCR buffer, 2.25 mmol L^-1^ MgCl_2_, 0.8 mmol L ^−1^ deoxyribonucleotide triphosphate (dNTP), 0.5 μmol L^-1^ of each primer, 2.5 U Taq DNA polymerase, and sterile filtered milli-Q water to a final volume of 50 μL. All reactions were carried out in a PTC-200 thermal cycler (MJ Research Co., New York, USA). PCR cycles included a 4 min initial denaturation at 94 °C, followed by 30 cycles of denaturation at 94 °C for 1 min, annealing at 53 °C for 30 s, extension at 72 °C for 1 min, and a 5-min final elongation step at 72 °C. PCR products were quality-screened and purified using the Qiangen Gel Extraction kit (Qiagen, Hilden, Germany).

### 2.5. 454 pyrosequencing and sequencing processing

Pyrosequencing was performed on a Roche Genome Sequencer FLX+ using Titanium chemistry by Macrogen (Roche Applied Science, Mannheim, Germany). Three standard flow-gram format (SFF) files were generated by 454 pyrosequencing. The SFF file was analyzed by the software package mothur (version 1.33.2) following the protocol provided by https://mothur.org/wiki/454_SOP. Briefly, De-noising and chimera analysis conducted with the AmpliconNoise (Quince et al., 2011) and UCHIME algorithms were used to reduce sequence errors. Furthermore, quality trimming was conducted to remove unwanted sequences shorter than 200 bp and reads containing ambiguous bases and with homopolymers longer than 8 bases. Remaining sequences were used to identify unique sequences by aligning with the SILVA-based bacteria reference alignment. Within unique sequences, the Uchime tool was applied to remove chimeras. Next, “Chloroplast”, “Mitochondria”, or “unknown” were identified and removed from the dataset. Subsequently, after calculating the pairwise distance and generating the distance matrix, a 97% identity threshold was used to cluster sequences into Operational Taxonomic Units (OTUs) according to the UCLUST algorithm (Edgar, 2010). For each OTU, the SILVA database was applied to annotate taxonomic information.

### 2.6. Data availability

Sequencing data are available in the NCBI SRA data repository under the project No. PRJNA679995.

### 2.7. Network construction and analysis

In order to determine the effects of different stand ages on microbiome associations in soils, underlying co-occurring bacterial taxa were depicted through co-occurrence network analysis. We divided all soil samples from all tea stands into four groups according to their land use types and stand ages: (1) Forest (F); (2) 3-20 year-old (Y3-20); (3) 40-50 year-old (Y40-50); and (4) 90 year-old (Y90). In order to reduce the complexity of the network, a Spearman’s correlation between two families was considered statistically robust if the Spearman’s correlation coefficient (r) was >0.8 and the p-value was <0.01 (Barberan et al., 2012). Meanwhile, a multiple testing correction using the Benjaminie Hochberg (FDR) method was applied to adjust the *p* values and reduce the chance of obtaining false-positive results (Benjamini and Hochberg, 1995). All robust correlations identified from pairwise comparison of family abundance form a correlation network in which the node represented bacterial family taxa and the edge represented a strong and significant correlation between the nodes. In addititon, we also generated sub-networks for each soil sample from meta-community networks by preserving OTUs presented in each tea stand with the subgraph function in igraph packages (Ma et al., 2016). To describe the complex pattern of interrelationship between bacterial taxa, a set of topological characteristics (number of nodes and edges, average path length, network diameter, average degree, graph density, clustering coefficient, and modularity) was determined using psych (Revelle, 2017), vegan (Oksanen et al., 2010) and igraph (Csardi and Nepusz, 2006) packages in R environment (Version No.: 3.60). Networks were visualized using the interactive platform Gephi (Bastian et al., 2009). In addition, 10,000 Erdos-Reyni (ER) random networks were generated to compare with the topology of real network with a random graph which connects each pair of nodes with any probability (Erdős and Rényi, 1960).

### 2.8. Statistical analysis

All statistical analyses were performed in R environment (Version No.: 3.60). To compare differences in soil physicochemical properties and alpha diversity of soil bacterial community between different stand ages, a repeated measures ANOVA, followed by multiple pairwise comparison using Tukey’s test at α = 0.05 was performed by ggpubr package. Linear discriminant analysis effect size (LEfSe) was performed to elaborate potential bacterial biomarkers (from phylum to genus) within soil microbiomes that specifically enrich different stand ages of tea plantations, based on *p* < 0.05 and a LDA score > 2.0 (Segata et al., 2011). Permutation multivariate analysis of variance (PERMANOVA) was employed to assess the effects of stand ages and sites on soil bacterial community using Adonis function of vegan package (Bell et al., 2014). Redundancy analysis (RDA) was conducted to identify soil physicochemical properties with significant impact on soil bacterial communities across different stand ages of tea plantations. Parameters that significantly explained variation in bacterial community were identified using forward selection (ordistep function of vegan package) with P value < 0.05. FAPROTAX (Functional Annotation of Prokaryotic Taxa) was applied to predict the microbial ecological function profiles by using the trans_func class of microeco package (Louca et al., 2016; Liu et al., 2021)).Non-metric multidimensional scaling (NMDS) analysis based on Bray-Curtis distances was used to evaluate the composition changes in microbial function groups between different sites and stand ages.

Maps of sampling sites were generated using GenGIS and soil physicochemical properties were ordinated by principal component analysis (PCA). The α – diversity (richness, evenness, and diversity) of soil bacterial communities was estimated based on OTUs. All indices were calculated using Vegan package in R environment. To assess changes in soil bacterial community structures among different tea managements and stand ages, principal coordinates analysis (PCoA) was used to calculate the gradient in compositional changes of bacterial microbial community (based on Bray-Curtis distances). The Spearman’s correlation analysis between the Euclidean distances of standardized sub-network topological parameters was applied to explore the effects of soil properties and stand ages on sub-network topological features. Soil properties (except for pH) were normalized before the correlation analysis.

## 3. Results

### 3.1. Soil physicochemical properties

Soil pH and 10 other soil properties (TOC, TON, C/N, MBC, MBN, AP, Exch.K, Exch. Ca, Exch. Mg and Exch. Na) were listed in Table 1. Based on detected soil properties, PCA showed that soil samples from the TRI site feature significant environmental heterogeneities across different stand ages (Fig. 1A) The content of TOC and TON increased along with the stand ages across the three sampling sites (Table 1), and the correlation analysis indicated that alongside the increased stand age of tea plantations, TOC and MBN in soil increased significantly (*p* < 0.01) (Fig. 1B and C). However, no consecutive changes along with increased stand age were recorded for other soil properties (e.g. MBC, AP, and Exch. K) (Table 1).

**Fig. 1.**
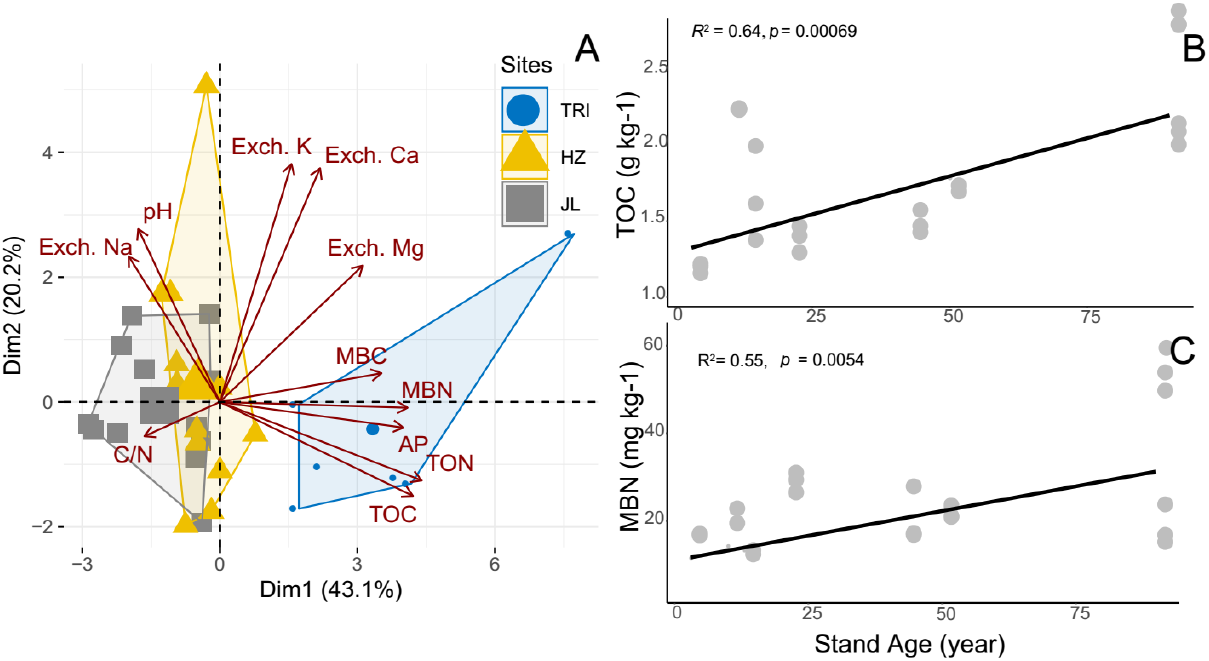
(A) Principal component analysis (PCA) based on soil physicochemical properties as variables. The sampling tea stands from same site were circulated and grouped as same color. (B) and (C) significant linear regression (*p* < 0.01) between total carbon (TOC)/ microbial biological nitrogen (MBN) in the soil and the stand age across all tea plantations.

**Table 1.**
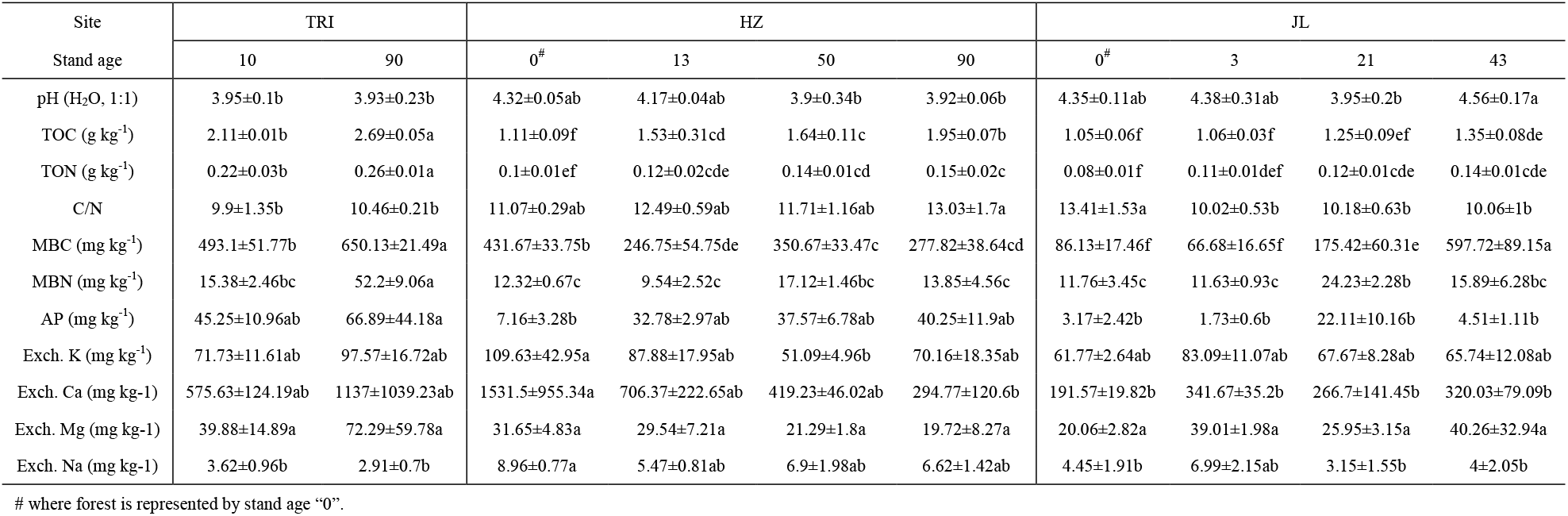
Soil physicochemical properties in tea plantations under different sites and stand ages

### 3.2. The bacterial component at different phylogenic level

Sequencing the amplicon libraries resulted in a total of 341,915 raw reads prior to quality checking and assigning reads to their respective sample. Average read length (± standard deviation) of reads before processing was 405 ± 96 bp. After quality trimming and assigning reads to different samples, 204,723 high quality reads remained in the dataset with an average length of 207 ± 4 bp.

The dominant bacterial phyla across all samples were Proteobacteria, Actinobacteria, Acidobacteria, Chloroflexi, and Firmicutes, while on average 15% of the reads could not be classified (Fig. 2A). To detect soil bacteria taxa that were significantly affected by tea stand age and location, the LEfSe analysis based on OTUs was applied to compare the differences. In general, the change trend of bacterial taxa varied from sites. LEfSe analysis revealed that 16 biomarkers affiliating with three phyla increased significantly (*p* < 0.05, LDA > 2.0) at the 90-year-old tea plantation, while 13 biomarkers within four phyla decreased significantly at the 10-year-old tea plantation at the TRI site. Moreover, LEfSe analysis demonstrated that few bacterial taxa increased with years of tea planting in JL and HZ sites compared to with forest soil (F).

**Fig. 2.**
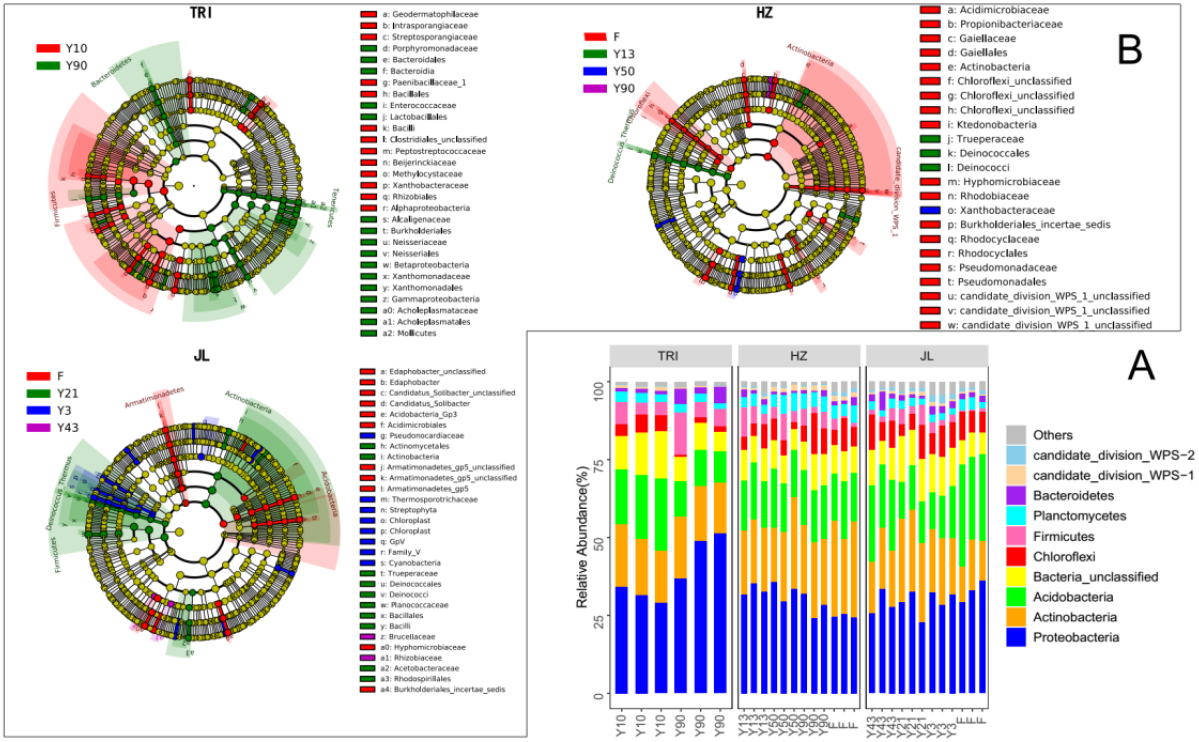
(A) Taxonomic structure of the soil bacterial microbiota at the phylum level. Only the 10 phylum with the highest mean relative abundance were shown, while the other phylum groups were grouped into “others”. (B) The effects of varying stand ages on the relative abundance of soil bacterial lineages were assessed through LDA Effect Size (LEfSe) with an absolute logarithmic LDA score threshold of 2.0 at three sampling sites (TRI, HZ and JL respectively). There are six circular rings in the cladogram, each circular ring deposit all taxa within a taxonomic level; the circular ring from inside to outside represents supergroup, phylum, class, order, family, and genus respectively

### 3.3. Bacterial community diversity and structure

The bacterial richness, or alpha diversity, varied widely across different sites (Fig. 3A). Soil bacterial richness did not change significantly at different stand ages across all sites except at the HZ site that saw a significantly higher richness in F than that in tea plantations. Further Spearman’s correlation analysis revealed that soil bacterial richness was significantly correlated with pH (R^2^ = 0.71, *p* < 0.001) and negative-correlated with TOC (R^2^ = −0.53, *p* < 0.001) in tea plantations (Fig. 3B and 3C, respectively).

**Fig. 3.**
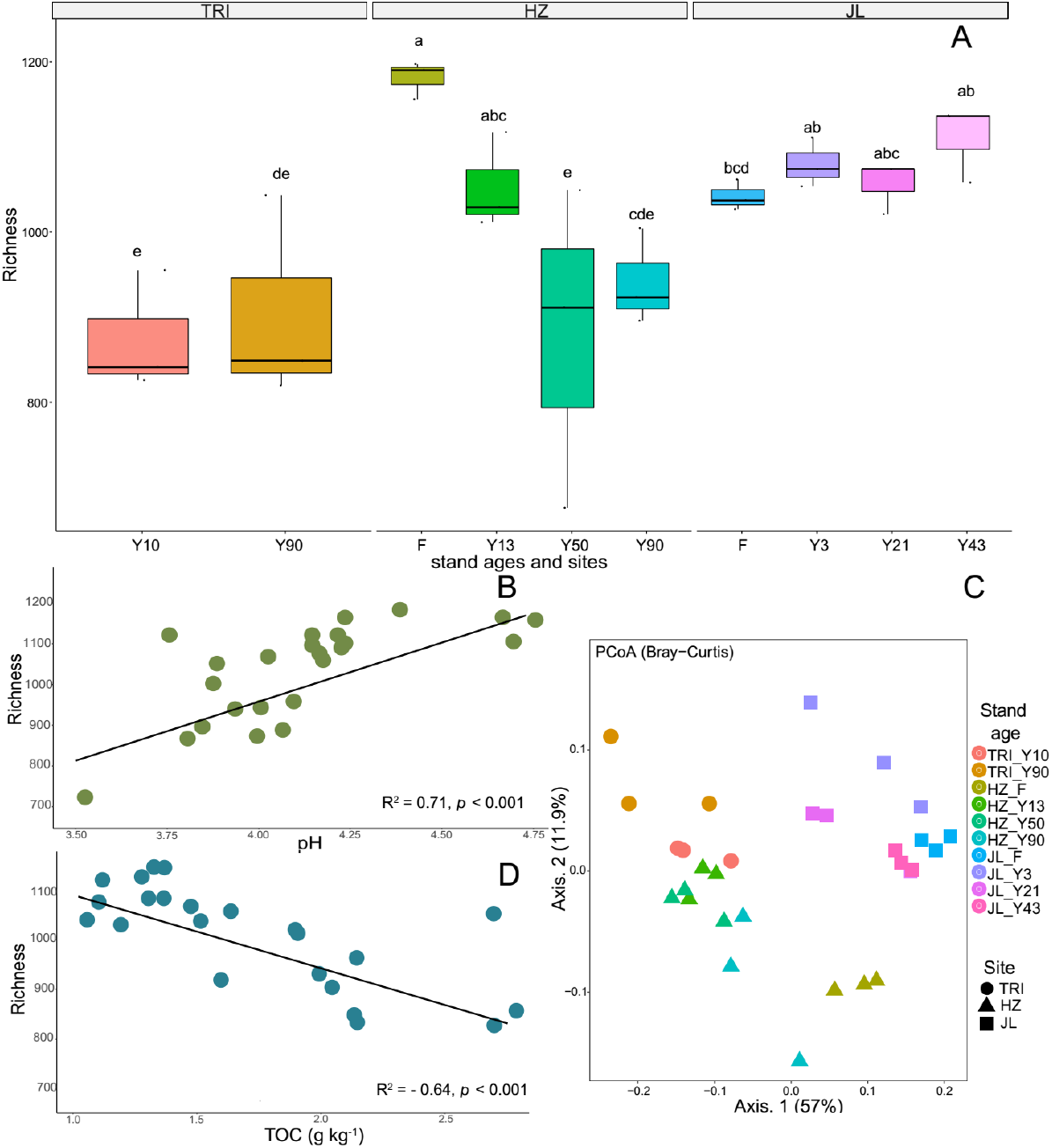
(A) The richness of soil bacterial community of tea plantation at varying stand ages and different sampling sites. (B) and (C) significant linear regression (p < 0.001) relationships between total carbon (TOC)/ soil pH in the soil and the richness of soil bacterial communities across all tea stands. (D) Principal coordinate analysis (based on Bray-Curtis distances) of soil bacterial community composition across varying stand ages and different sampling sites. The samples separated by sites (TRI, HZ, and JL, respectively; represented by different shape) and stand ages (F (adjacent forest); represented by different colors).

To explore how changes in microbiome structure and composition correlated with sampling sites and stand age, we computed the between-sample diversity (β-diversity) using Bray-Curtis distance. Axis 1 and axis 2 explained 57% and 11.9% of the total variation in bacterial community structure, respectively Principal coordinate analysis (PCoA) of bacteria community structure revealed in Fig. 3D that soil samples from different sites with different stand ages were generally clustered separately.

### 3.5. Soil bacterial functions

A total of 22 functional sub-categories (relative abundance > 1%) within 5 major categories “Energy source”, “C-cycle”, “N-cycle”, “S-cycle” and “Others” were identified and linked to the microbial communities across different tea stand ages and sites (Fig. 4A). No consecutive change of these functional sub-categories was observed alongside with the increasing tea stand years in all three sites. Nevertheless, the results in Fig. 4A showed that in TRI site, longer stand year of tea plantation soils had a higher relative abundance of chemoheterotrophy, photoheterotrophy, fermentation, cellulolysis, chitinolysis, nitrate reduction, nitrate respiration, nitrogen fixation and aerobic ammonia oxidation. In addition, in JL site, the relative abundance of dark hydrogen oxidation increased with the stand ages of tea plantation. When converting forest to tea plantation, in HZ site, the relative abundance of most C-cycle functions decreased. Non-metric multidimensional scaling (NMDS, based on Bray-Curtis distance) plot of all the 22 sub-categories showed the separate clusters of functional categories between TRI and other two sites soils (Fig.S2A). In TRI and JL sites, the clusters from different tea stand soils separated from each other, suggesting that there were significant differences in the function of soil microbial communities between different tea stand ages (Fig. S2B).

**Fig. 4.**
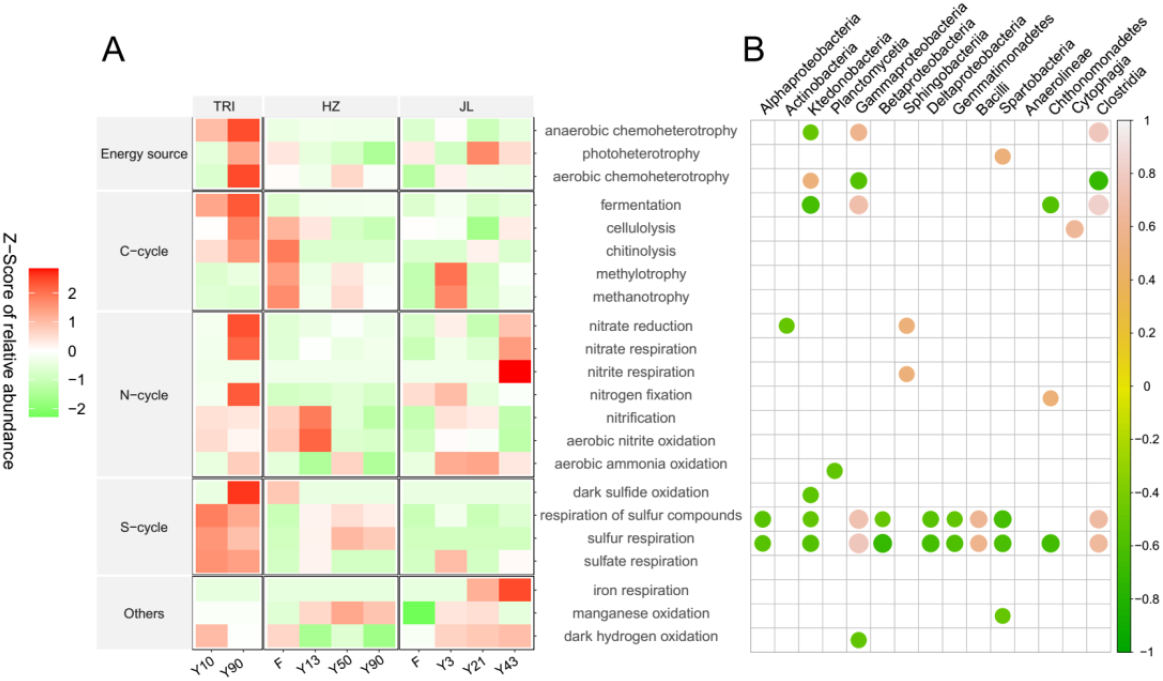
Function predictions of microbial communities in tea plantation soils across varying stand ages and different sampling sites by FAPROTAX. The relative abundance of each functional categories is normalized and represented by Z-score. (A) Comparison of dominant functional categories; (B) The Spearman’s correlation between 15 most abundant microbial classes and predicted functional categories.

Spearman rank-order correlations detected the significant correlations (*p < 0.05*) between dominant microbial classes (the top 15 most abundant classes) and functional categories, suggesting that most microbial classes possessed the functions of respiration of sulfur compounds, and sulfur respiration (Fig. 4B). Additionally, the classes of Ktedonobacteria, Gammaproteobacteria and Clostridia were significantly correlated with the functions of anaerobic chemoheterotrophy, aerobic chemoheterotrophy and fermentation (*p* < 0.05). For N-cycle category, the classes of Actinobacteria, Sphingobacteria, Chthonomonadetes showed the significant correlations with different sub-categories (Fig. 4B).

### 3.6. Co-occurrence network analysis of soil bacterial community

For co-occurrence network analysis, we divided all samples into four groups: (1) Forest (F); (2) 10-20 year-old (Y10-20); (3) 40-50 year-old (Y40-50); and (4) 90 year-old (Y90). Subsequently, four networks (F, Y3-20, Y40-50, Y90) were constructed to test the effect of stand age on soil bacterial communities’ association. Overall, nodes in all networks were assigned to 15 bacteria phyla and three unclassified groups. Co-occurrence networks were markedly different among different stand ages (Fig. 5A). Most links were derived from the phyla of Proteobacteria, Actinobacteria, Acidobacteria, Firmicutes and Bacteroidetes across the 4 network we constructed (Fig.5B). However, the proportion of each phyla varied with different networks. In the F network, three phyla (Proteobacteria, Actinobacteria, and Acidobacteria) accounted for over 75% of the total links, but this proportion decreased in other tea stand networks (Fig. 5B). We then investigated the correlations between key topological parameters of the subnetworks and stand ages by Spearman’s correlation analysis. Average path length (APL) and centralization betweenness positively and significantly correlated with stand age (*p* < 0.05), suggesting that the importance of individual bacterial community groups became more uniform as stand age increased. However, the number of edges and nodes showed no significant correlation with stand ages (Fig. 5C). Further Spearman’s correlation analysis between topological parameters and soil physicochemical properties revealed that soil C, N and P were all significantly correlated with some key parameters of subnetworks (e.g. APL, diameter and centralization betweenness) (Fig. 5D). When comparing the network parameters we calculated in Fig. 5E between the four co-occurrence networks, the result showed that the number of positive edges was much higher than that of the negative edges across the soils from all stand ages as well as that of forest soil. Furthermore, values relating to APL, clustering coefficient, and numbers of clusters in those empirical networks of various tea plantations and forest were higher than those of their respective, identically sized Erdose-Reyni random networks. This indicates that the empirical networks had significant “small-world” modularity and hierarchy of their topological properties (Fig. 5E). Further structural analysis showed that the clustering coefficient and edge numbers of the networks increased along with stand ages (Fig. 5E), indicating that the increased tea plantation stand age made soil bacterial community associations more complex and tightened. In addition, compared with F soil, all soil bacterial communities of tea plantations had a relatively high APL value (Fig. 5E).

**Fig. 5.**
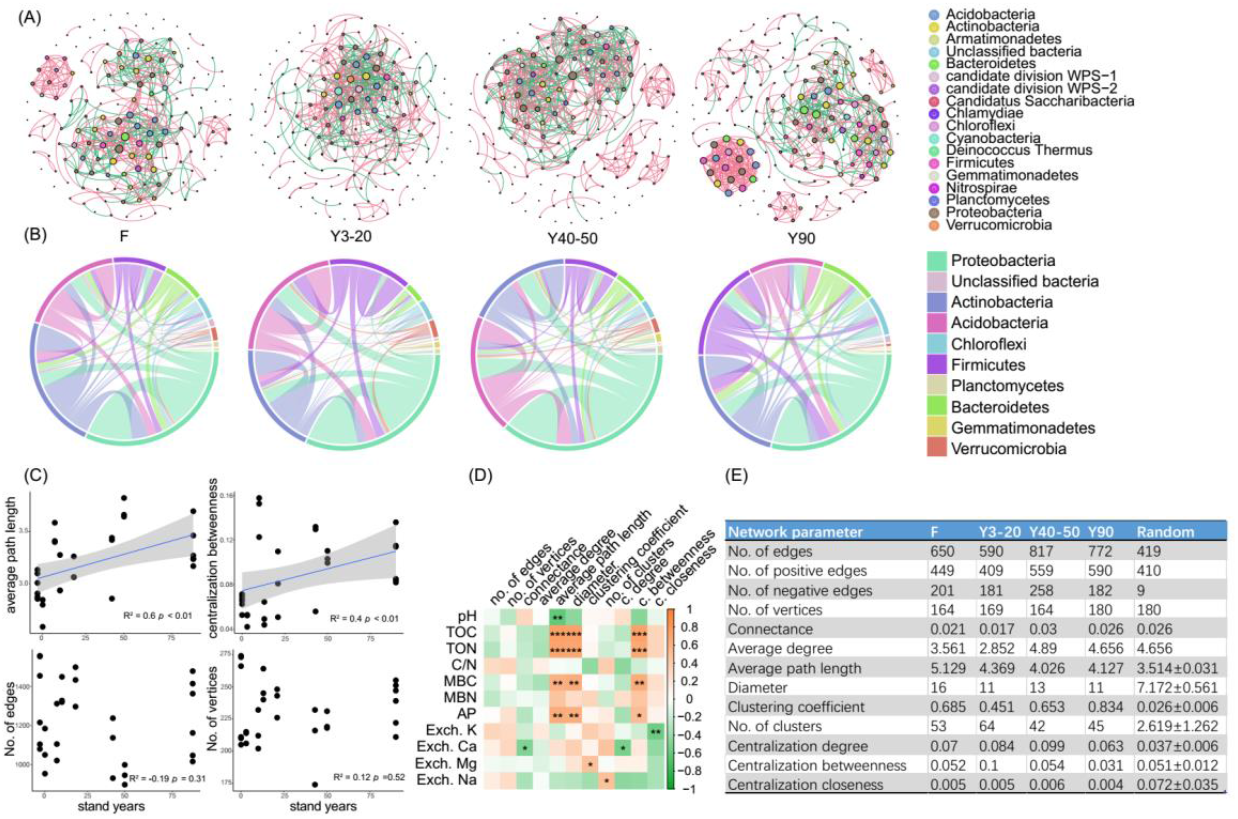
(A) The co-occurrence networks visualize the effects of varying stand ages of tea plantations (Y3-20, Y40-50, Y90) and the adjacent forest (F)) on the co-occurrence pattern between soil bacterial taxa at family level The node size is proportional to the abundance of taxa, and the nodes represent bacterial taxa at family level. The edges are colored according to interaction types; positive correlations are labeled with green and negative correlations are colored in pink. (B) CIRCOS plots showing the distribution of links among the top 10 interacting phyla in networks Y3-20, Y40-50, Y90 and F. (C) Linear regression relationships between tea stand ages and key topological parameters of all subnetworks (average path length, centralization betweenness, No. of edges and No. of verticles). (D) Spearman’s correlations between soil physicochemical properties and topological parameters in all subnetworks. Significant correlations are marked by * (*p* < 0.05), **(*p* < 0.01), and ***(*p* < 0.001). (E) Topological parameters of the networks Y3-20, Y40-50, Y90 and F.

### 3.4. Relationship of soil properties with the structure, function and co-occurrence partten of soil bacterial communities

Redundancy analysis (RDA) was applied to study the effects of soil properties on the structure of soil bacterial communities based on OTU abundance. The ordination diagram showed that bacterial community change was significantly correlated with soil variables: TOC, TON, MBC, pH, and AP&K (*p* < 0.05, Monte Carlo test) (Fig. 6A). The first two axes of RDA can explain 41.5% and 12.5% of the total variation. In addition, both sampling sites and stand age can significantly affect soil bacterial communities (PERMANOVA test, *p* <0.01), and stand age (R^2^ = 0.569) outcompeted sampling sites (R^2^ = 0.377; Fig. 6B) for controlling bacterial community composition. In addition, to identify the edaphic drivers of soil microbial communities in tea plantation, we correlated the composition of taxonomic and functional communities and topological parameters with soil properties. The mantel correlations showed that soil C and N as well as AP were the strongest correlates of both taxonomic and functional composition (Fig. 6C). At the same time, TOC, TON, MBC and MBN were also strongly self-correlated. For the co-occurrence network, soil Exch. K content was the significant correlate (*p* < 0.05) (Fig. 6C).

**Fig. 6.**
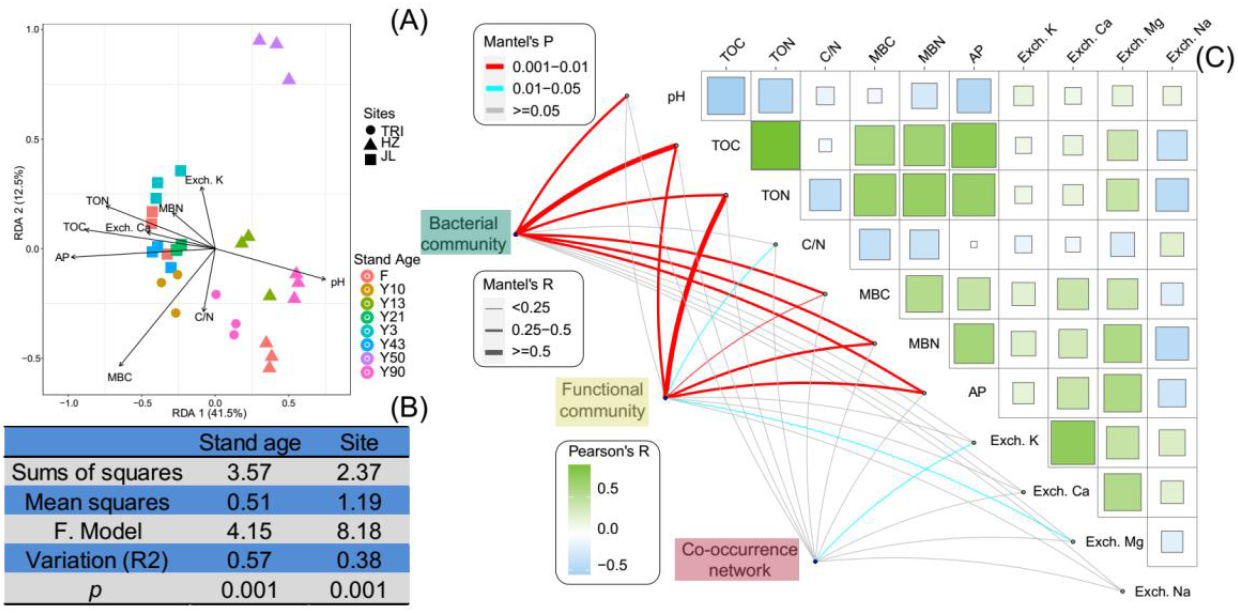
(A) Redundancy analysis (RDA) of the relationships between soil physicochemical properties and bacterial communities under different sites and stand ages. The samples separated by sites (TRI, HZ, and JL, respectively; represented by different shape) and stand ages (including the adjacent forest (F); represented by different colors). (B) The effects of stand age and sampling site on soil bacterial communities of tea plantations based on permutation multivariate analysis of variance (PERMANOVA). (C) Pairwise comparisons of soil physicochemical properties are shown, with a color gradient denoting Pearson’s correlation coefficients. Bacterial (based on the relative abundance of OTUs) and functional (based on the relative abundance of functional categories predicted by FAPROTAX) community composition and co-occurrence network (based on the topological parameters of all subnetworks) are related to each soil properties by Mantel tests. Edge width corresponds to the Mantel’s R statistic for the corresponding distance correlations, and edge color denotes the statistical significance based on 9,999 permutations.

## 4. Discussion

Soil microbial communities are of particular relevance in tea cultivation, since soil microbiota demonstrate reservoirs of microorganisms colonizing tea plantations, and contributing to improved yield and tea quality (Mortimer et al., 2015). To characterize the effects of long-term monoculture and other environment factors like location and soil properties on soil bacterial communities, we investigated soil bacterial communities of tea plantation at different stand ages and associated forests across three different sites in eastern China.

### 4.1. Effect of soil properties on shaping soil bacterial communities

Our study revealed that soil TOC and MBN in tea plantation are significantly positively correlated with stand age (Fig. 1B&C). This is in line with previous studies reporting the plantation age as a critical factor affecting SOC and N dynamics during land use change, in particular on tea plantations (Pansombat et al., 1997; Wang et al., 2016). This demonstrates that long-term tea plantations result in a significant accumulation of organic C and N. The increase in the amount of soil micro-aggregates in long-term tea plantations can be a reason for such an increase in TOC and MBN, as micro-aggregates are the most predominant pools of SOC and other nutrients (Wang et al., 2016). In addition, the application of long-term organic and mineral fertilizers into tea plantations could also result in the accumulation of organic C and N in the soil.

Soil pH is an important factor affecting soil bacterial communities in regional and global scales, which has been confirmed in other ecosystems (Delgado-Baquerizo et al., 2018). For instance, Zhou et al. (2017) found that richness of soil bacteria was significantly negatively correlated with soil pH in rubber plantations (pH: 3.94-4.41) but in an oil-contaminated soil, pH was positively associated with bacterial diversity (pH: 7.49 - 9.20) (Jiao et al., 2016). Our research has shown that in an acidic tea plantation, soil pH was strongly positively correlated with the alpha diversity of soil bacterial communities (Fig. 3B). This finding is in line with previous studies (Griffiths et al., 2011), which have illustrated that soil pH is positively correlated with bacterial alpha diversity, and alpha diversity of bacterial communities was highest at a near-neutral pH.

In addition to the importance of some commonly accepted variables like C, N, and soil pH in shaping soil microbial communities, our study also supports the finding that soil base cations like Ca and K are important in shaping the composition and co-occurrence network of the bacterial communities of tea plantation (Fig. 6). These base cations act as nutrients or structural components of the cells of living microorganisms (Tripler et al., 2006). When considering the drivers of soil bacterial communities, the availability of Ca and K mostly impact the bacteria involved in the dissolution of soil minerals (e.g., mineral-weathering bacteria) (Puente et al., 2004). The effects of Ca and K on structure of soil bacterial communities have been reported in forest soils (Uroz et al., 2011) and agricultural soils (Schmidt et al., 2019). Our study reaffirmed the importance of these base cations in shaping bacterial community structures and intra-taxa associations in tea plantation soils.

### 4.2. Effects of tea plantation stand age on microbial communities

The alpha diversity of soil bacterial communities remained stable during long-term tea plantations (Fig. 3A). This sort of stability was also observed in a 20 years tea plantation (Li et al., 2016). Zhao et al. (2012) proposed that in a monoculture cropping system like tea plantation, rhizosphere effects are the critical factor that determine bacterial community diversity, and the toxicity and accumulation of antimicrobial substances due to long-term cropping as well as the specific acidic soil environment which may suppress the development of bacterial populations. One or combination of these factors could explain the stability of alpha diversity observed in our study. Furthermore, soil properties like pH are known to intimately determine bacterial diversity and community composition (Bissett et al., 2011). As shown in Table 1, soil pH did not show a significant response to long-term tea plantations at each site, which could also result in the insignificant change of the alpha diversity. In addition, despite the acidic soil environment, the increasing TOC and MBN with successive tea planting may also contribute to the stability of the alpha diversity, as soil organic matter and nutrients have a profound effect on microbial diversity (Montecchia et al., 2015). Previous studies (Jangid et al., 2008; Lee-Cruz et al., 2013) have shown that the alpha diversity of soil bacterial communities can be declined by the conversion of forests to long-term agriculture management. In contrast, our finding at the JL site did not agree with this conclusion, and we suggest here that after forests were converted to monoculture agricultural system, it does not necessarily mean that the bacterial community’s diversity is reduced or lost. In addition, our study also revealed that some key soil functions related to C and N cycles shifted after converting forest to tea plantation (Fig. 4A). This disagreement is mostly because the effect of cultivation on alpha diversity and soil functions strongly depends on the nature of the soil and cultivation type (Coller et al., 2019). Since the quality of SOM was already low at pH 4.3 under forest, 0.4 unit decrease in pH over long-time plantation didn’t make a considerable difference in SOM quality. Therefore, the microbial response remained comparable between tea plantations and forests.

Considering the between-samples variability, both PERMANOVA and PCoA analyses indicated that sampling location and stand age can significantly affect the beta diversity of soil bacterial communities (*p* <0.01), and stand age (R^2^ = 0.569) more significantly affected the beta diversity than sampling location (R^2^ = 0.377; Table 2). It has been previously suggested that geographical origin is the dominant factor in determining the structure of soil bacterial communities in vineyards (Coller et al., 2019), which is partially consistent with our finding in tea plantation soils. Importantly, we found that stand age could indeed shape the structure of bacterial communities in tea plantation soil. Because this change in microbial community structure was mostly induced by C and N increases with tea stand age, the bacterial community structure is mainly affected by environmental variables (Jiao et al., 2016). In general, our study confirmed that environmental variability caused by long-term monoculture and spatial variability (tea plantation sites) determined the structure of bacterial communities in tea plantation soil.

### 4.3. Interactions among soil bacteria taxa were strengthened by long-term tea monoculture

Referring to the studied bacterial communities, the 16S rRNA sequencing indicated that Proteobacteria, Actinobacteria, and Acidobacteria were the dominant taxa across all samples in tea plantation soils. This finding is consistent with previous Chinese tea plantation soil studies (Li et al., 2016). Co-occurrence network analysis across all stand ages also revealed that most of the nodes belonged to these three phyla. In addition, we found that the tea plantation networks were nonrandom and typically matched the topological features of a small-world and intrinsic modular architecture (Barberan et al., 2012). This typical “small-world” characteristic in tea plantation soils made the networks more strengthened than random associations (Watts and Strogatz, 1998).

Interestingly, to our knowledge, this study is among the first reporting that long-term tea monoculture tightened soil microbial associations. One possible explanation is that changes of some taxa were sensitive to C and N increase induced by tea planting. The LEfSe analysis detected several carbon- or nitrogen-susceptible taxa in the Rhizobiales, Xanthomonadales, and Burkholderiales, which have previously been reported as keystone taxa in agricultural ecosystems linked to C and N metabolism in soils (Li et al., 2015). In our study, the function prediction also revealed that some bacterial taxa like Actinobacteria, Chthonomonadetes were correlated with the key processes in C and N cycles (Fig. 4B). Another explanation is that greater nutrient availability like C and N in the soil could subsequently strengthen microbial interactions in order to enhance the efficiency of resource turnover that benefits tea growth (Shi et al., 2016; Zhao et al., 2019). Lastly, according to the topological characteristic analysis, long-term tea monoculture and agropedogenesis reduced betweeness centralization and the links of key bacterial taxa in the networks, which could partially contribute to the tightening of soil bacterial associations in tea plantation (Kuzyakov and Zamanian, 2019).

In order to display the chronosequence development of networks with stand ages, we merged tea stands soils from different sampling sites into one stand age group. Despite that, we acknowledge that this merge could obscure the effects of geographical parameters on network construction, but nonetheless this study is the first to demonstrate that long-term monoculture has tightened soil microbiome network associations in tea plantations.

## 5. Conclusions

Through comparison of three independent sites, we found that long-term tea monoculture led to significant increases in TOC and MBN. This C and N availability improvement in tea plantation soils could contribute to tea yield growth and be of greater benefit to tea monocultural systems. The analysis of 16S rRNA genes of bacterial communities revealed that the structures of soil bacterial communities were significantly changed by the stand age of tea plantations, sampling locations, and land-use conversion. Stand age outcompeted sampling locations for controlling the composition of bacterial communities. Interestingly, this study is the first of its kind to report that long-term tea monoculture tightened soil microbiome associations through co-ocurrence network analysis.

## Acknowledgements

This research was supported by the National Key R&D Program of China (2017YFE0107500) and the Research Fund for International Young Scientists of National Natural Science Foundation of China to K.Z. (Grant number: 42050410320). Heng Gui would like to thank the support from the National Natural Science Foundation of China (NSFC Grant 32001296) and Yunnan Fundamental Research Projects (grant NO. 2019FB063).

