## Supplementary materials for "Variations in soil nutrient dynamics and bacterial communities in long-term tea monoculture production systems"

**Supplementary Fig. S1** Sampling sites of tea plantations in Zhejiang Province, China
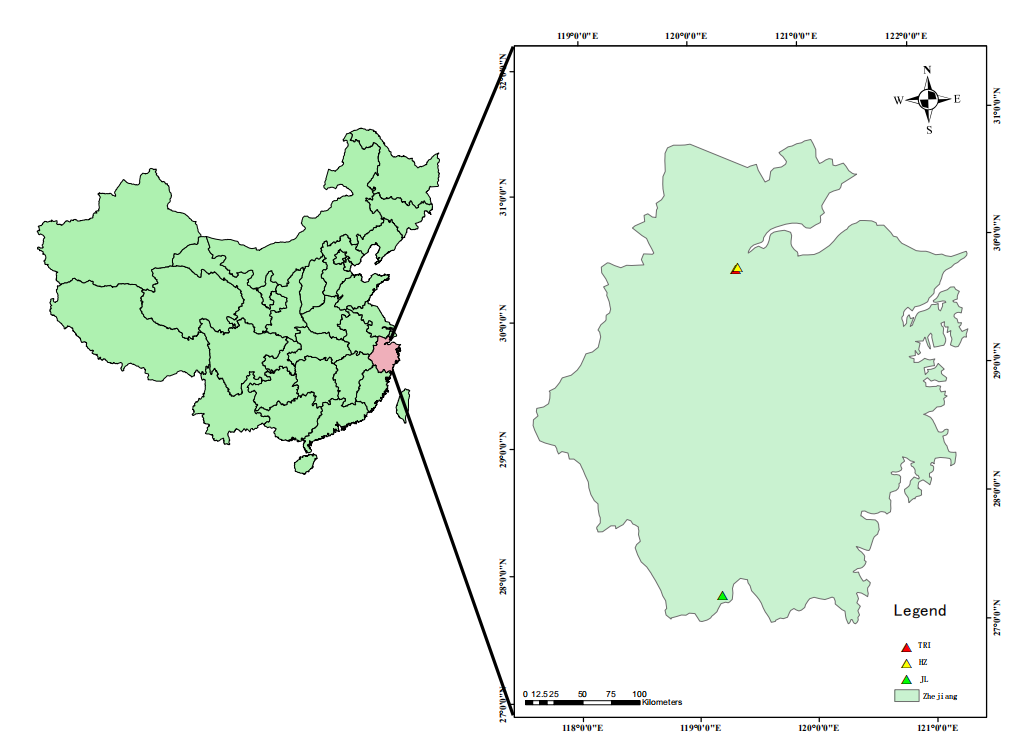
(TRI: the Tea Research Institute of the Chinese Academy of Agricultural Sciences; HZ: Wenjiashan village, Hangzhou city; JL: Jingning county, Lishui city).

**Supplementary Table S1.** Selected information for the three tea plantations located in Zhejiang province, China

| Sites | Annual Mean Temperature (℃) | Annual mean precipitation (mm) | | US Soil classification | |
| --- | --- | --- | --- | --- | --- |
| TRI | 17 | | 1533 | | Ultisols |
| HZ | 17 | | 1533 | | Ultisols |
| JL | 17.5 | | 1599 | | Ultisols |

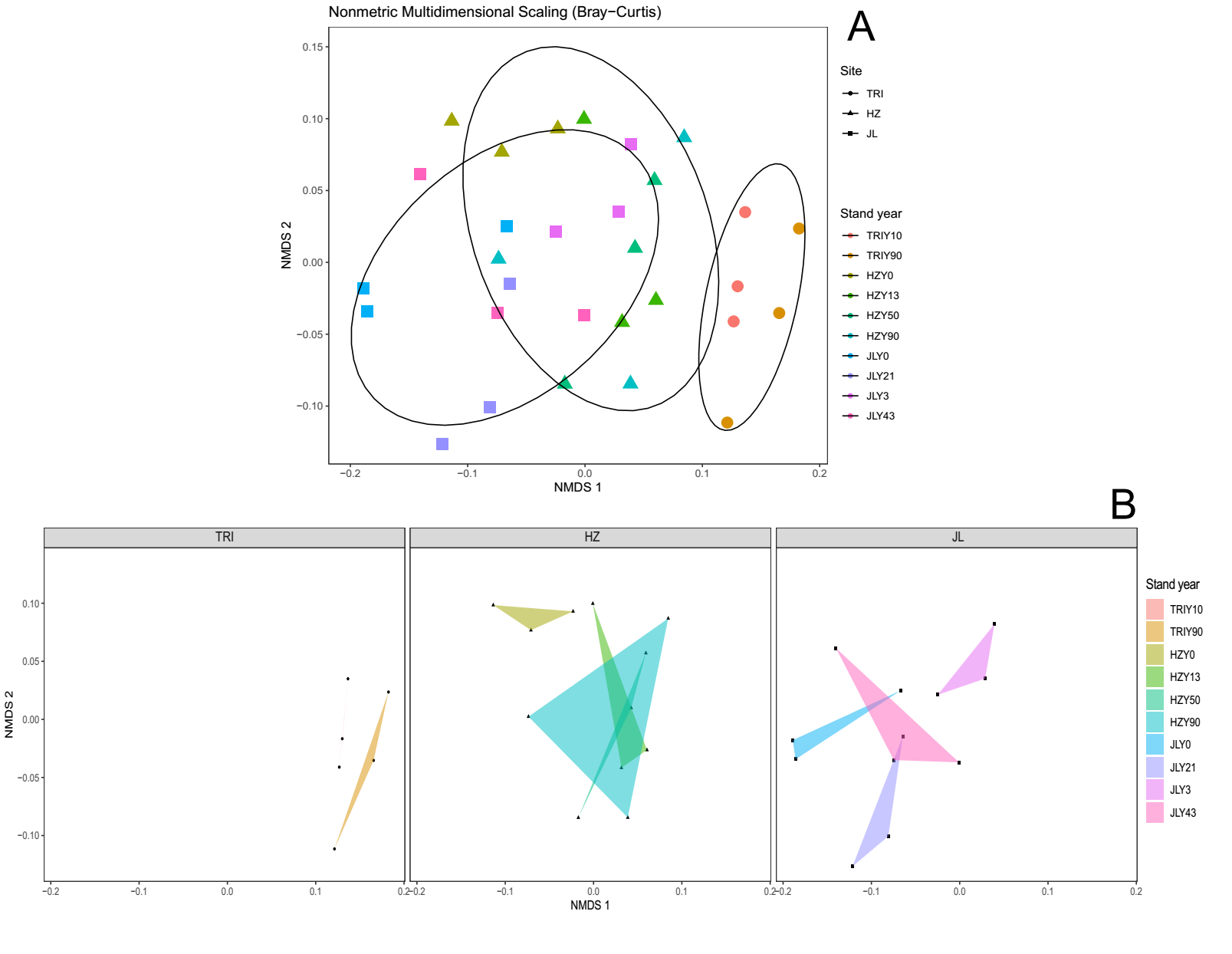

**Supplementary Fig. S2** Non-metric multidimensional scaling (NMDS) ordination based on Bray-Curtis distances under functional categories predicted by FAPROTAX (Stress value = 0.188). (A) and (B) show the NMDS ordination of all 3 sites and 3 each site separately. The samples separated by sites (TRI, HZ, and JL, respectively; represented by different shape) and stand ages (F (adjacent forest); represented by different colors).
